# Junctional and cytoplasmatic contributions in wound healing

**DOI:** 10.1101/2020.02.21.959379

**Authors:** Payman Mosaffa, Robert J. Tetley, Antonio Rodríguez-Ferran, Yanlan Mao, José J. Muñoz

## Abstract

Wound healing is characterised by the re-epitheliation of a tissue through the activation of contractile forces concentrated mainly at the wound edge. While the formation of an actin purse string has been identified as one of the main mechanisms, far less is known about the effects of the viscoelastic properties of the surrounding cells, and the different contribution of the junctional and cytoplasmic contractilities.

In this paper we simulate the wound healing process, resorting to a hybrid vertex model that includes cell boundary and cytoplasmatic contractilities explicitly, together with a differentiated viscoelastic rheology based on an adaptive rest-length. From experimental measurements of the recoil and closure phases of wounds in the *Drosophila* wing disc epithelium, we fit tissue viscoelastic properties. We then analyse in terms of closure rate and energy requirements the contributions of junctional and cytoplasmatic contractilities.

Our results suggest that reduction of junctional stiffness rather than cytoplasmatic stiffness has a more pronounced effect on shortening closure times, and that intercalation rate has a minor effect on the stored energy, but contributes significantly to shortening the healing process, mostly in the later stages.

**Author summary:** We simulate the wound healing process of epithelia in the absence of substrate. By analysing the recoil process we are able to fit the viscoelastic properties of the monolayer, and study the influence of contractility at junctions and at the interior polymer network. We numerically simulate the whole wound opening and closure process, and inspect which mechanism has a more pronounced effect in terms of energy barrier and wound closure rate. We conclude that while junctional stiffness seems to be more effective than bulk stiffness at speeding up the closure, the increase of intercalation process is the mechanism with the lowest energy cost.

## Introduction

Wound healing is a fundamental process for maintaining integrity and functionality in epithelia, tissues, and organisms [1]. The most accepted mechanisms for wound closure are Rac-dependent crawling using cell protrusions [2], and the formation of a contractile supra-cellular actomyosin cable at the wound edge (purse-string mechanism) [3]. However, in embryonic and larval tissues, either the elastic substrate is not present or no crawling has been observed. In these cases, the first mechanism is absent and closure relies mostly on the actomyosin cable and cell-cell interactions [4], which is the focus of our study in this article.

Extensive analyses have reported the key mechanical factors affecting wound closure. The effects of myosin heterogeneity [5, 6], purse-string assembly rate [6], or tissue fluidity [4] have been studied. However, far less is known about the contribution of junctional and cell medial contractility and stiffness during closure, and the energy barrier that they may impose during the intercalation process. We here simulate wounding of the *Drosophila* imaginal disc epithelium [4], and numerically analyse the influence of junctional versus medial cortex stiffness and contractility in a vertex model. Their differential roles during tissue remodelling have been already reported [7], and thus we expect that their different properties will influence cell polarisation and intercalation that takes place during wound healing.

The modelling of wound healing on substrates using vertex models has been presented for instance in [6, 8–11], while homogenised continuum models can be found in [12–14], which allow the inclusion of myosin and calcium dynamics [13, 15]. Due to the importance of junctional mechanics, and the absence of a force contribution between cells and their underlying substrate in our reference *ex vivo* system, we will employ a customised vertex model with no crawling. However, both medial and junctional cortices are differentiated in conjunction with a viscoelastic rheological model.

The tailored vertex model and the comparison with the experimental rate of tissue recoil will enable us to calibrate the material viscoelastic properties. The employed rheological model resorts to a Maxwell-like viscous model where the rest-length is able to dynamically adapt [16]. Similar rest-length based models have been recently adopted when modelling embryonic tissues [17, 18] or stress relaxation in suspended monolayers [19]. We will apply these ideas to the wound healing process.

We will assume uniform constant viscoelastic material properties, as it have been measured experimentally [5], and uniform ring contractility at the wound edge. Although is has been experimentally observed that the supra-cellular actin cable is not uniformly continuous [5, 10], we will coarsen its net effect at the wound front.

Our numerical model incorporates some differences with respect to our previous hybrid model [9], mainly on the treatment of the wound edge and wound interior. We first highlight these differences, and then discuss our results and compare them to relevant experimental outputs. We place special emphasis on the role of the intercalation and remodelling process, as recently studied in [4]. We additionally inspect the role of the intercalation in terms of contributions from the cell junctions and medial forces across cell boundaries, and show that they have unequal effects on the evolution of the stored energy and the closure time.

## Materials and Methods

### Hybrid Vertex Model

The proposed computational approach is based on a discrete description of the tissue forming a flat monolayer. The cell boundaries are described through a set of connected vertices ***y***^***I***^ forming the *vertex network*, as is customary in vertex models [20]. In addition, we also include the *cell centres network*, defined by a set of nodes ***x*^*i*^**, *i* ∈ {1, 2, …, *N*_*nodes*_}, with *N*_*nodes*_ the number of cells in the tissue, and a triangulation that defines the connectivity between cells, i.e. cells *i* and *j* are connected if a bar element *ij* is present in the triangulation.

In order to obtain the connectivities between cells, we use a *Graded* Delaunay triangulation (GDT). It corresponds to a modification of the Delaunay algorithm, where the aspect ratio of triangles is not necessarily optimal. Instead, connectivity between two adjacent triangles is swapped only if the aspect ratio of one of the triangles is larger than a parameter *δ*. More specifically, given two triangles 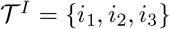 and 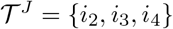, with nodal positions 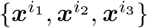 and 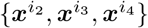, the connectivity is modified to 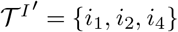 and 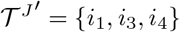 if

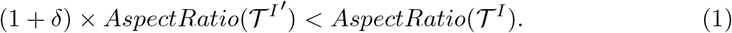

*AspectRatio* 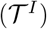 is measured as the ratio between the radius of the circumcircle and the incircle of triangle 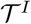. Condition in (1) is checked for *I* ∈ {1, 2, …, *N*_*tri*_}, with *N*_*tri*_ the number of triangles in the tissue. We point out that as a consequence of condition (1), the model will have a higher number of elongated cells as we increase parameter *δ*, and that for *δ* = 0 standard Delaunay triangulation is used. For this reason, we call this parameter *intercalation restriction*.

The topology and geometry of the cells centres can be defined by a matrix 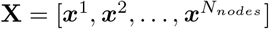 and a matrix 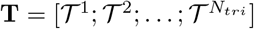 defining the connectivity of the cells. The positions of cell centres and vertices in the two networks are not independent. Vertices are in general located at the barycentres of each triangle. In other words, given a triangle 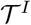, the coordinates of the associated vertex ***y*^*I*^** are given by,

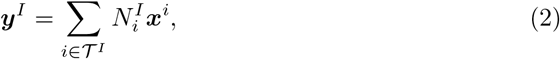

with *N*_*i*_ three interpolation weights, which we set to 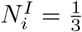, for *i* = 1,2,3 and all triangles *I* = 1, …, *N*_*tri*_. While the cell centre network eases the description of the connectivity changes, the vertex network allows us to include the cortex contractility, as will be explained later. We note that similar coupled interpolations have been used on other vertex models [21, 22].

### Curved cell boundaries at the tissue edge

Since the boundary of the boundary cells lies outside the triangular nodal network, vertices at the tissue boundary, denoted by ***y*_*b*_**, do not follow the interpolation in (2). Instead their location is initially defined from the vertices of triangles that have bar elements at the boundary of the triangulations, and subsequently computed through mechanical equilibrium. Appendix S1 describes how the positions of the boundary vertices are defined, preserving the initial area of cells, while mechanical equilibrium is described in the next section.

As a result, the whole tissue geometry and connectivity is given by matrices **X**, **T** and 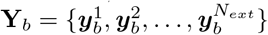, with *N*_*ext*_ the number of external vertices. Cell centre and vertex geometry, **X** and **Y**_***b***_, are computed through mechanical equilibrium, and connectivity **T** is recomputed at each new time step *t*_*n*+1_ from the resulting coordinates resorting to the GDT. Figure 1 illustrates the process of obtaining 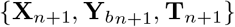 from 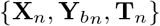.

**Fig 1.**
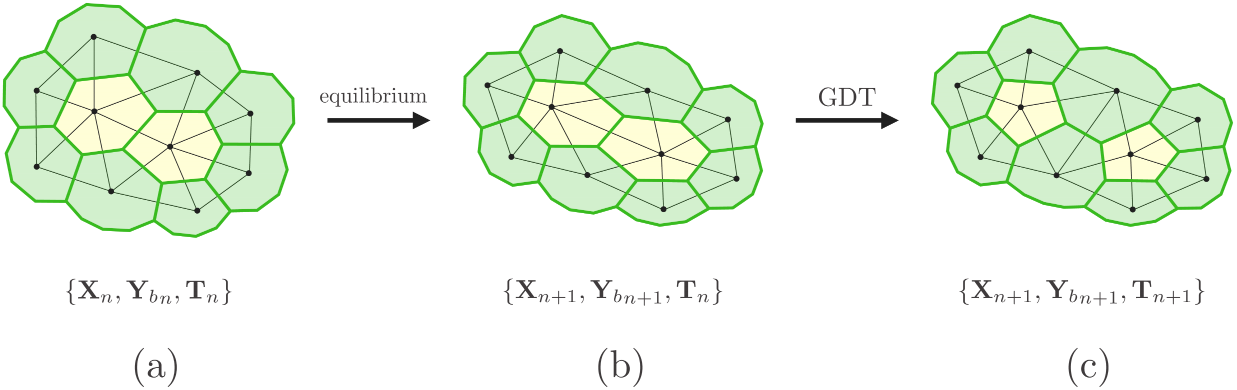
Schematic of the computational process for retrieving position of cells and cells connectivities 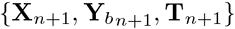 at time *t*_*n*+1_ from the same properties at time *t*_*n*_. (a)→(b): Computation of new nodal positions **X**_*n*+1_ and new positions of external vertices 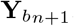. (b)→(c): Computation of cell connectivities by resorting to GDT of the new nodal positions

**Fig 2.**
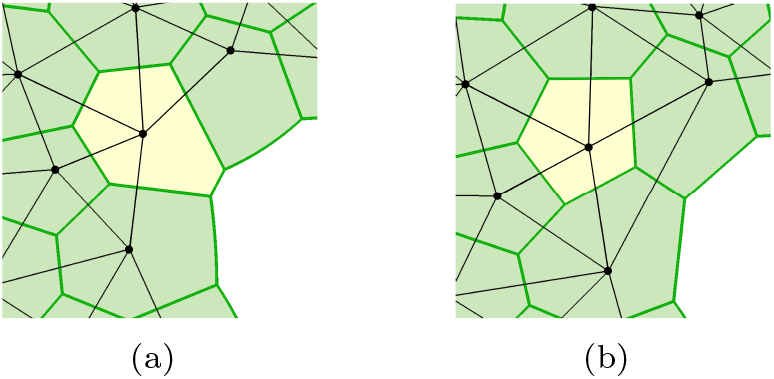
Schematic representation of the computation of cell connectivity at the tissue edge. (a) Cell, in yellow, contributing to the tissue boundary by a short edge (**T**_*n*_). (b) Cell, in yellow, does not contribute to the tissue boundary anymore, through computation of cells connectivity (**T**_*n*+1_).

### Cell connectivity at the tissue edge

Since the external boundary of the cells at the tissue edge is not defined directly through tessellation of the triangular nodal network, the cell connectivity at the tissue edge cannot be computed by resorting to the GDT. Instead, we employ another criterion which is based on the ratio between the length of the cell side at the tissue edge, *l*, and the cell perimeter, *P*. An intercalation is applied when this ratio is smaller than a certain threshold. The restriction on intercalation on the external cells then reads,

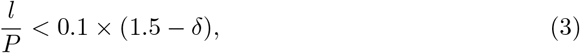

where the numerical values in 3 have been chosen so that the intercalations at the tissue edge resemble those in the bulk of the tissue, while being controlled by the same parameter *δ*.

### Mechanical equilibrium

#### Total elastic energy and equilibrium equations

Similar to other vertex models [6, 11, 20], our equilibrium equations are constructed by minimising a total potential function, which in our case is given by,

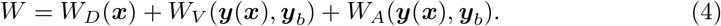

The terms *W*_*D*_(***x***) and *W*_*V*_ (***y***(***x***), ***y*_*b*_**) account respectively for the contribution of the bar elements in the cell centre and vertex networks. We also add the term *W*_*A*_(***y***(***x***), ***y*_*b*_**) which penalises area changes. The specific form of these contributions will be detailed below. As yet, the equations of mechanical equilibrium can be derived by minimising the total elastic energy *W* with respect to the system principal kinematic variables (degrees of freedom): nodal positions in **X** and the position of exterior vertices in **Y**_***b***_. This minimisation yields the following system of non-linear equations,

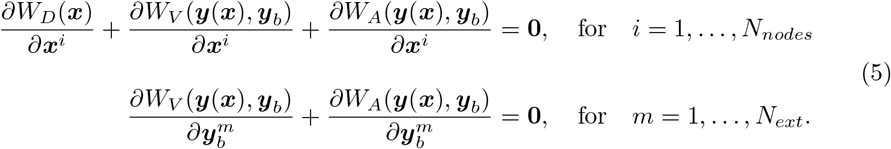

The derivatives given above can be obtained through standard mathematical manipulations. The equations are solved at each time-step through an iterative Newton-Raphson process, which requires full linearisation of the equations. The explicit expression of the equations and the iterative scheme can be found in [9].

### Nodal and vertex mechanics

Triangulation **T** of the tissue domain results in *N*_*D*_ pairs *ij* of connected nodes, defining a bar element each, and *N*_*V*_ pairs *IJ* of vertices. Each nodal bar element represents the forces between two cells, while vertex bar elements represent forces at cell junctions. Forces are deduced from the nodal and vertex energy function *W*_*D*_ and *W*_*V*_, which read,

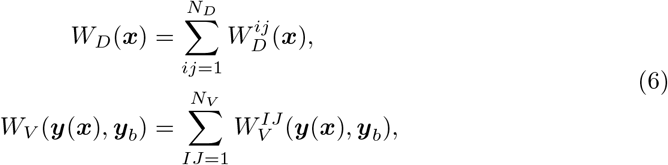

with the elemental functions given by

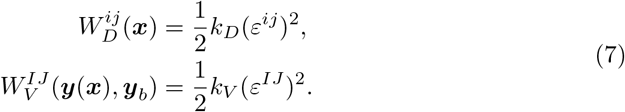

Material parameters *k*_*D*_ and *k*_*V*_ are respectively the intra-cellular and cell boundary stiffnesses, while 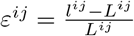 and 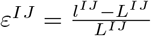 represent the nodal and vertex scalar elastic strain measures. Lengths *l*^*ij*^ = ||***x***^*i*^ − ***x***^*j*^|| and *L*^*ij*^ are the current and *reference lengths* of elements in nodal network, and *l*^*IJ*^ = ||***y*^*I*^** − ***y*^*J*^**|| and *L*^*IJ*^ represent the corresponding parameters in the vertex network.

### Area constraint

As the number and the size of the cells is considered approximately constant within the tissue, we consider a cell volume invariance in the model, which is applied here by imposing a two-dimensional area constraint through the energy term,

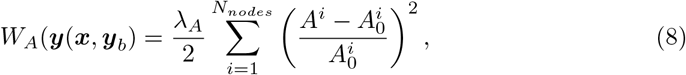

where *λ*_*A*_ is a penalisation coefficient and 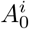 and *A*^*i*^ are the initial and the current areas of cell *i*, respectively. The value of *λ*_*A*_ is chosen such that relative area variations do not exceed 10%, as as experimentally reported in 3D measurements [5].

### Tissue Ablation

Wounding in the *Drosophila* wing disc is carried out by laser ablation of a number of cells within a region located at the interior of the tissue [4]. In the proposed computational approach, we simulate this by removing a number of cell centre nodes inside a circle with radius *r* and centred at the interior of the nodal network, which represents the area ablated in experiments. This will leave an empty area in the interior of the nodal network, as well as a new tissue edge (wound boundary), which needs to be constructed by new non-coupled boundary vertices 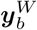.

Figure 3 depicts the wounding process in the vertex model. Due to the tissue elasticity, a recoil of the tissue and associated stretching of the boundary at the wound edge takes place.

**Fig 3.**
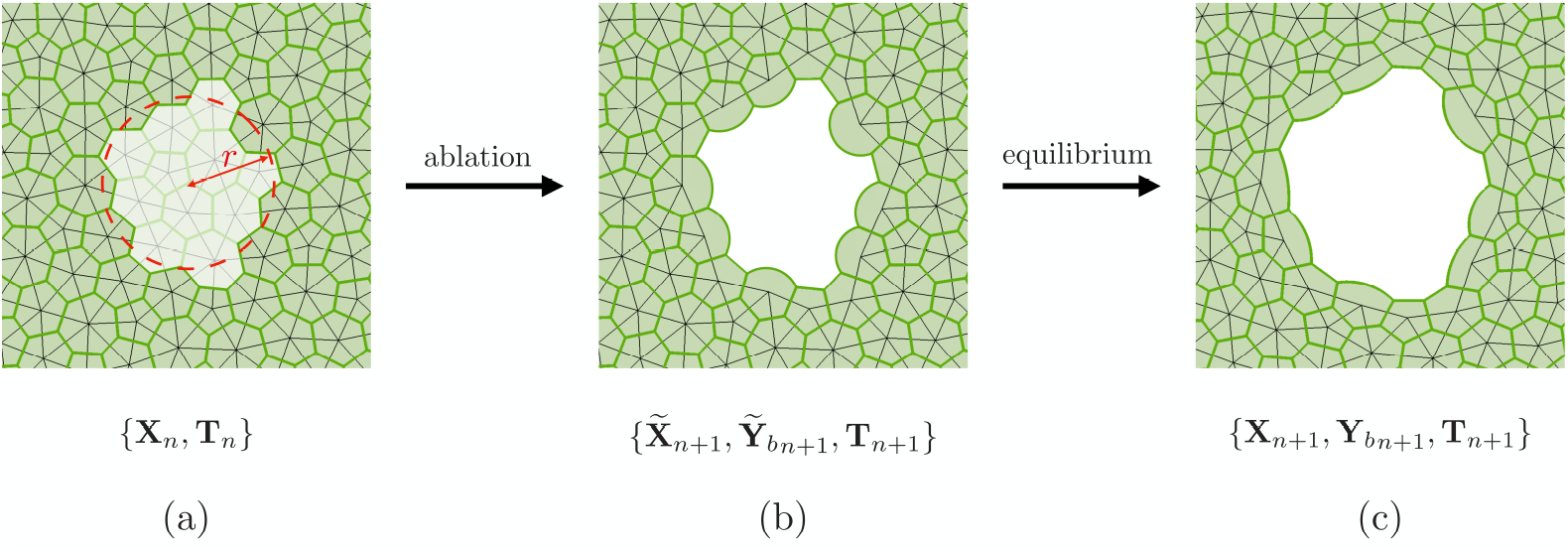
(a) Nodal and vertex configuration before ablation at time *t*_*n*_. (b) Ablation imposed by removing nodes located inside the laser beam with radius *r*, and construction of vertices 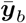 at the edge of the wound, preserving the current cells area at time *t*_*n*_. (c) New equilibrated cells configuration at time *t*_*n*+1_ computed from mechanical equilibrium.

### Rheological model

The strain measures in (7) are written in terms of the difference between the measurable current length *l* and the rest-length *L*, which is used as an internal variable, and is not necessarily constant. In order to mimic the viscoelastic response of cells and cell junctions [23–25], we resort to a dynamic law of the rest-length evolution

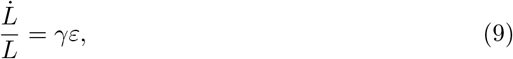

where *γ* is the *remodelling rate*, and *ε* is the elastic strain used in (7). It has been previously shown that such a rheological model reproduces Maxwell viscoelastic behaviour [16], and that it can be employed to simulate tissue fluidisation [19, 26] or cell cortex response in embryogenesis [17, 27].

Since cells exhibit inherent contractility exerted at the cortex cross-linked structure [28], we modify the evolution law in (9) as

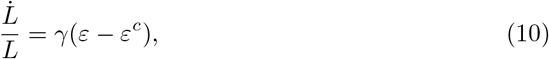

with *ε*^*c*^ a contractility parameter. Note that according to this law, the rest-length will remain constant whenever the elastic strain *ε* equals the value of the contractility *ε*^*c*^, which in each simulation takes a constant value.

The differential equation in (10) is discretised resorting to a Backward Euler scheme. The unknown rest-lengths *L*_*n*+1_ are thus written in terms of the current lengths *l*_*n*+1_ and inserted in the non-linear equations in (5), which are solved at each time-step *t*_*n*+1_, allowing the new nodal positions 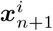 and boundary vertices 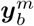 to be found.

Due to cell reorganisation, new bar elements may be created at each time-step. This requires the redefinition of the rest-lengths for some of the elements. We resort to an Equilibrium Preserving Mapping, which computes new elemental rest-lengths that minimise equilibrium errors at vertices and nodes. Further details of this mapping may be found in s [9]. The hybrid vertex model has been implemented in Matlab (R2018, The Mathworks, Inc.) and is available for its use (see online Supplemental Information).

We use different contractilities for the nodal and vertex networks, denoted respectively by *ε*_*D*_ and *ε*_*V*_. In addition, in order to mimic the traction of the actomyosin cable during wound closure, the edges at the wound front have a different contractility denoted by 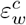. The values of these and other model parameters are commented on in the next section.

## Results and discussion

Wound closure is simulated by first applying cell ablation and subsequently applying a constant value of 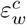. Figure 4 shows the time sequence of the process in the numerical simulation. The time lapse between wounding and the application of ring contractility, *t*_*c*_ − *t*_*ab*_, is fixed to 240 seconds, in agreement with *in vivo* experiments.

**Fig 4.**
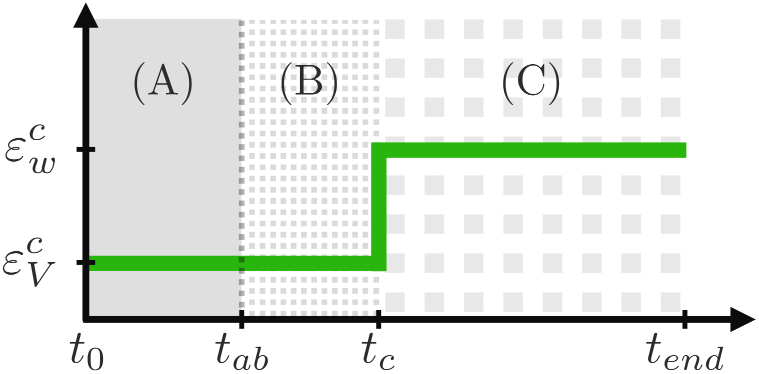
Time sequence of ablation.(A) Tissue preconditioning. During *t*_*ab*_ − *t*_0_ the tissue reaches a homeostatic contractile state. (B) Wound recoil. At *t*_*ab*_ ablation is applied by removing the cells within the wound area, after which the wound edge undergoes a recoil during *t*_*c*_ − *t*_*ab*_ = 240 seconds to recover a new equilibrated configuration. (C) Wound closure. At *t*_*c*_ the contractility at the wound front edge increases from 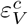 to 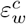 in order to mimic the traction of the actomyosin cable.

Our numerical model depends on eight material and numerical parameters: the material stiffnesses *k*_*D*_ and *k*_*V*_, the remodelling rates *γ*_*D*_ and *γ*_*V*_, the GDT intercalation restriction, *δ*, and the contractilities *ε*^*c*^, *ε*^*c*^ and 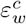.

We use the time evolution of the wounded area *A*(*t*) in order to fit these parameters. Indeed, the recoil process allows us to set some reference values of material constants (*k*, *γ*, *ε*)_*D*_ and (*k*, *γ*, *ε*)_*V*_, while the closure rate is mostly determined by the values of 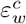 and *δ*, in conjunction also with (*γ*_*D*_, *γ*_*V*_). In Appendix S2 we explain how each factor affects the resulting curve *A*(*t*) and how the fitting is accomplished. Table 1 gives the obtained reference values of the parameters, and in Figure 5 we plot the evolution of *A*(*t*) for the experimental and numerical results using the reference values in Table 1.

**Table 1.**
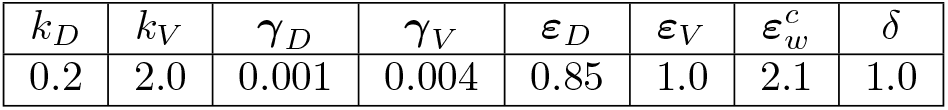
Reference model parameters fitting the experimental results.

**Fig 5.**
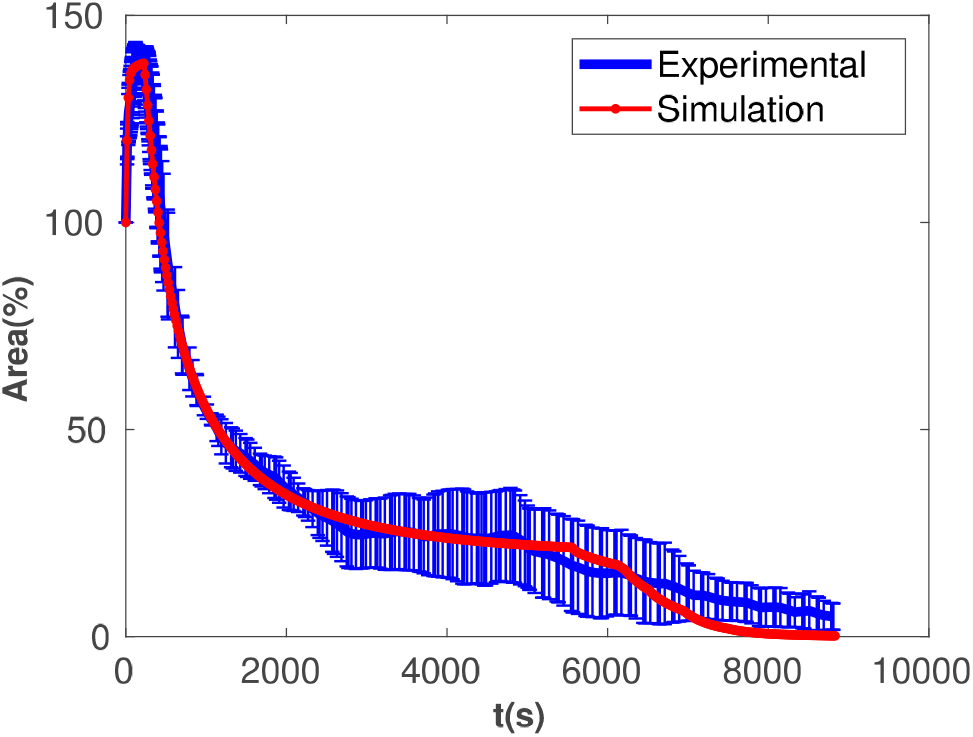
Experimental and fitted simulation results. Vertical blue bars indicate standard deviation from mean values using *N* = 5 measurements.

Four snapshots of two representative simulations are given in Figure 6. From the comparison of the wound evolutions it can be verified that different combinations of *k*_*D*_ and *k*_*V*_ yield different stresses at the cell boundaries, while keeping similar values of the area. In those cases where the fitted values of stiffnesses *ε*_*D*_ and *ε*_*V*_ at the nodal and vertex network are not unique, we determine their values from the trend of *A*(*t*) during closure. Our numerical tests revealed that increasing values of contractility 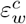 and lower values of tolerance *δ* yield higher rate of diminishing area and a larger number of intercalations, reducing the closure time. Conversely, larger values of tolerance *δ* and lower values of 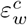 promotes the formation of wound with many surrounding cells, yielding rosette-like wound shapes.

**Fig 6.**
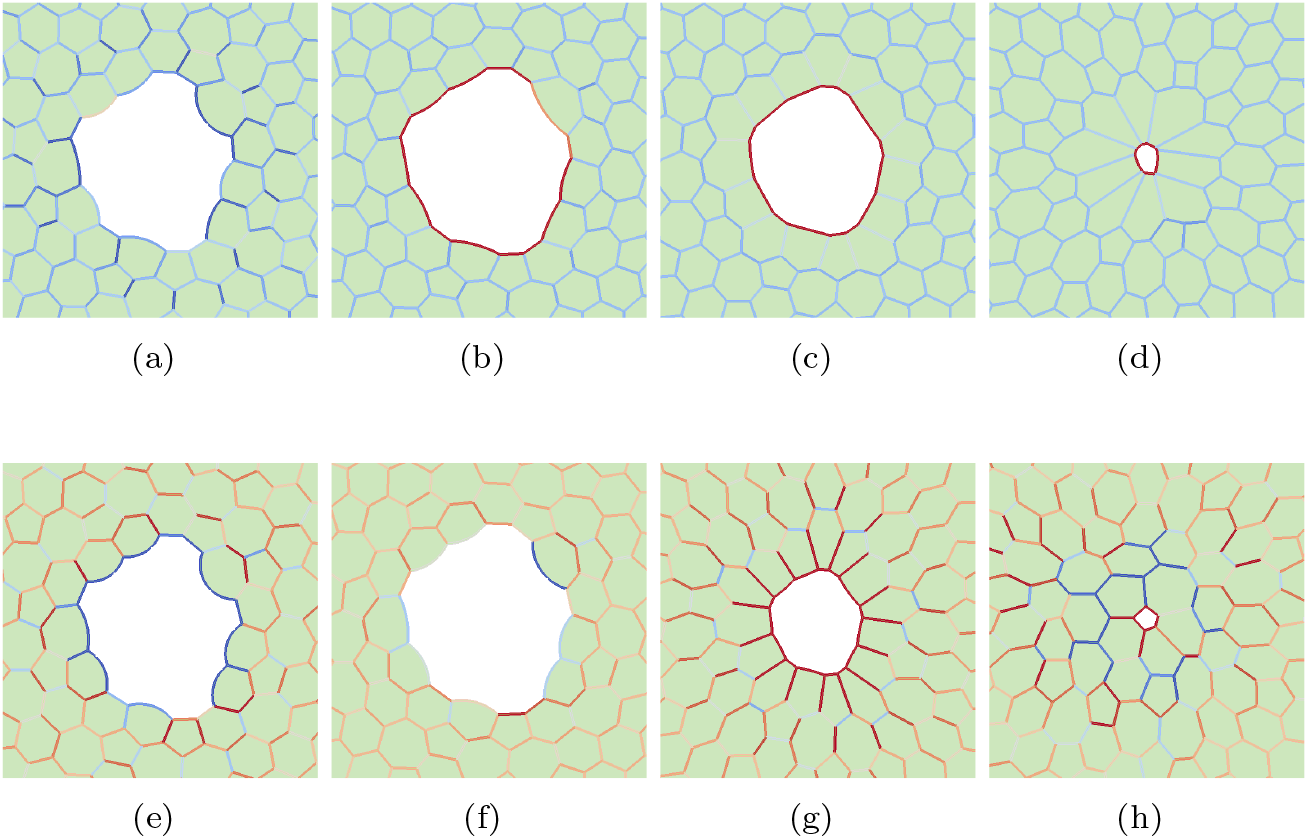
Snapshots of wound closure for two sets of material parameters. Red level indicates level of tension at edges. (a)-(d): 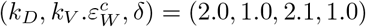. (e)-(h): 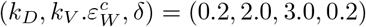. Times correspond to (a),(e): just after ablation, (b)-(f): at maximum recoil, (c)-(g): *t* = 10 min, and just before full closure, (d): *t* = 75 min, (h): *t* = 21 min.

It is worth pointing out that the presence of intercalation events at the wound edge gives rise to a disruption of the cell orientation towards the wound edge [29], allowing cells shape to relax towards a more isotropic distribution, despite imposing a purse string mechanism. Furthermore, although we impose a strain driven closure (rest-length at wound edge is such that the elastic strains converge towards the target value 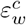), the total stress and strain are not homogeneous, as also experimentally measured [5].

With the aim of gaining further insight into the mechanisms that drive and prevent wound healing and that may increase closure rate, we also plot the time evolution of the elastic energy *W*_*E*_ in Figures 7 and 8. The first figure plots energy evolutions with a constant value of *δ* and different values of 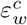, while the second plots the opposite combination of parameters. In particular, we note that the former indicates that lower purse string contractilities impose lower elastic energies but at the expense of longer closure times. The plot in Figure 8 instead confirms that by allowing more elongated cells, full closure takes a longer time, but with a similar final elastic energy.

**Fig 7.**
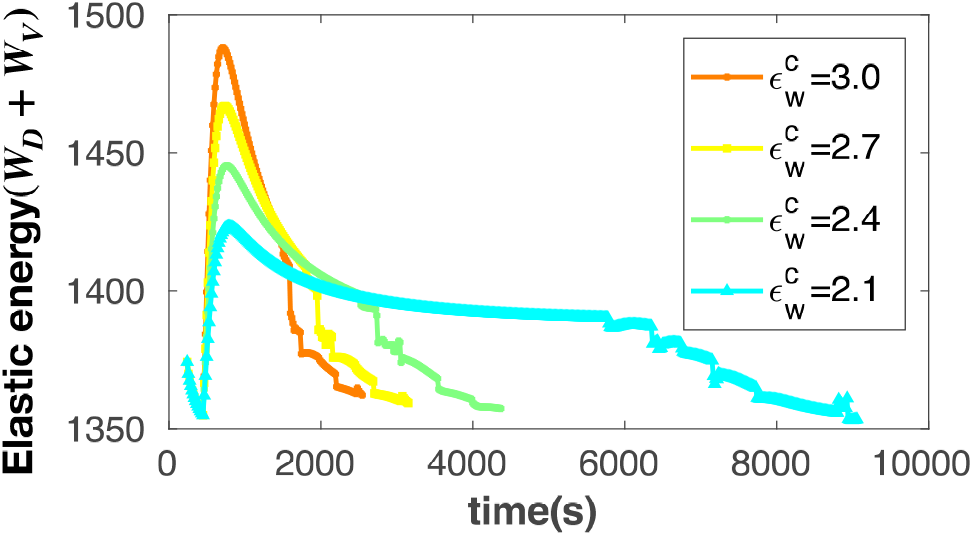
Time evolution of elastic energy, *W*_*E*_, for *δ* = 1.0 and a range of 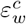.

**Fig 8.**
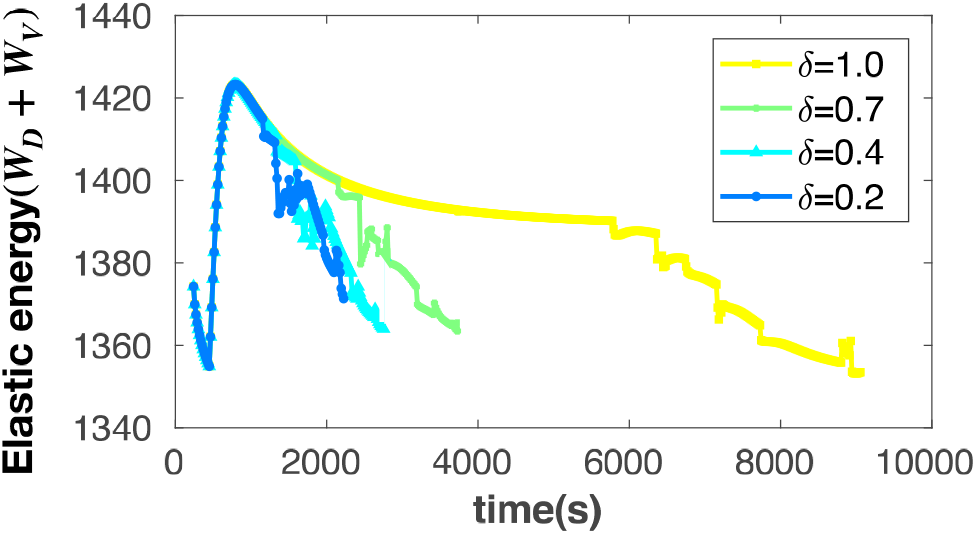
Time evolution of elastic energy, *W*_*E*_, for 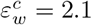 and a range of *δ*.

In both plots the faster the wound closes, the higher the final stored energy, although energy differences are relatively small. Also, we point out that energy levels at closure are always lower than energy before ablation, but higher than at maximum recoil. The fact that far fewer junctions are present during ablation and intercalation, may explain this energy drop with respect to initial state before wounding. However, the application of 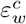 implies an energy supply, which may be biologically explained by actin flow, or higher consumption of ATP by myosin motors at the wound edge.

From the elastic energy (total energy minus the area contribution, *W*_*E*_ = *W* − *W*_*A*_), and area evolution in figures 7 and 8, we also construct phase diagrams where the elastic energy W_E_ is plotted against relative area *A*/*A*_*0*_. Figures 9 and 10 show the associated phase diagrams. The former indicates that higher wound tractions impose much higher energy peaks, which take place at decreasing values of relative area. Interestingly, the intercalation restriction has no effect on the energy evolution. Indeed, all the curves in Figure 10 collapse almost on a single energy-area curve, suggesting that the presence of intercalation, despite minor fluctuations, has no justification in terms of energy terms, but may be able to even reduce the closure time by approximately %75.

**Fig 9.**
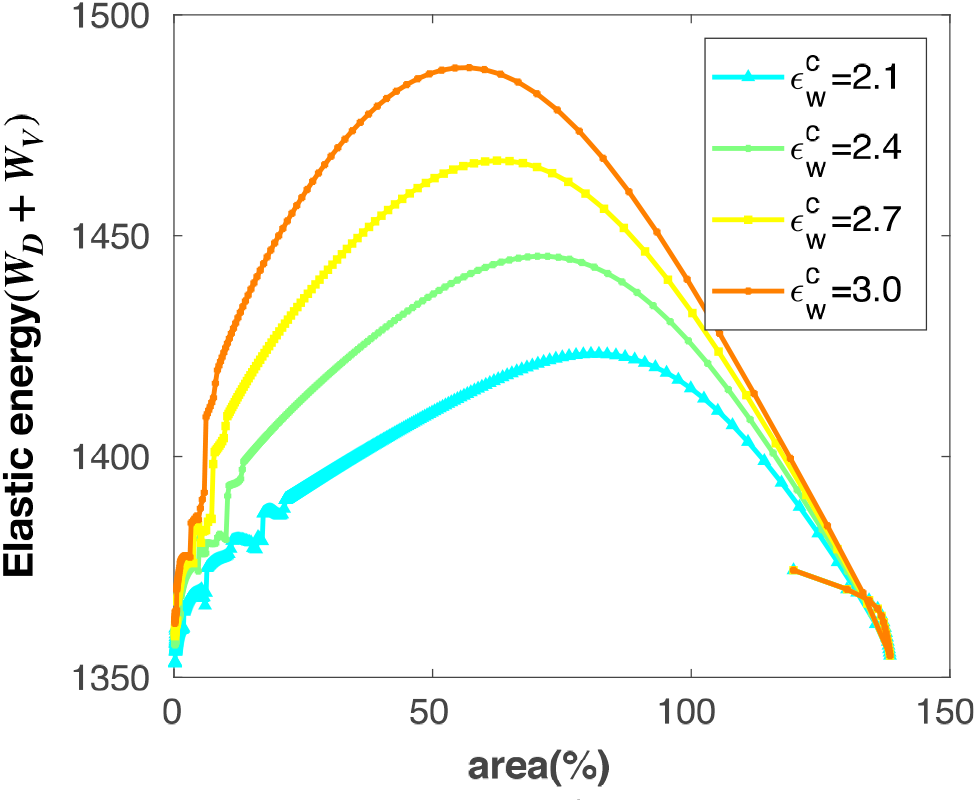
Elastic energy versus relative area for *δ* = 1.0 and a range of 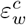.

**Fig 10.**
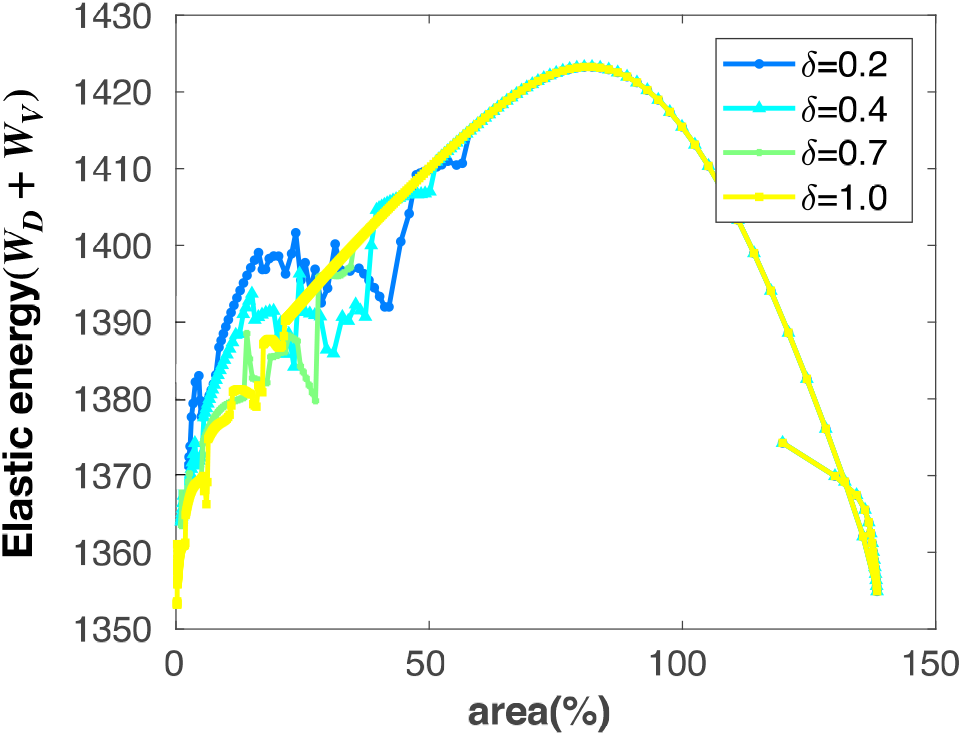
Elastic energy versus relative area for 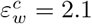 and a range of *δ*.

The vertex model presented here has the ability to distinguish between junctional and non-junctional cortex contributions. For this reason, we have analysed the relative effect of nodal and vertex stiffness parameters, *k*_*D*_ and *k*_*V*_. Figure (11) indicates that although increasing values of both stiffnesses lead to longer closure times, the increase of vertex stiffness has a more pronounced effect on the total time. When inspecting the energy profiles in Figure 12, we also observe that too large values of *k*_*V*_ may prevent full closure due to the wound contraction being insufficient to surmount the required energy barrier. Also, for sufficiently low values of *k*_*V*_, the energy level after closure approximates the one after the recoil phase, although a slight energy difference remains in all cases.

**Fig 11.**
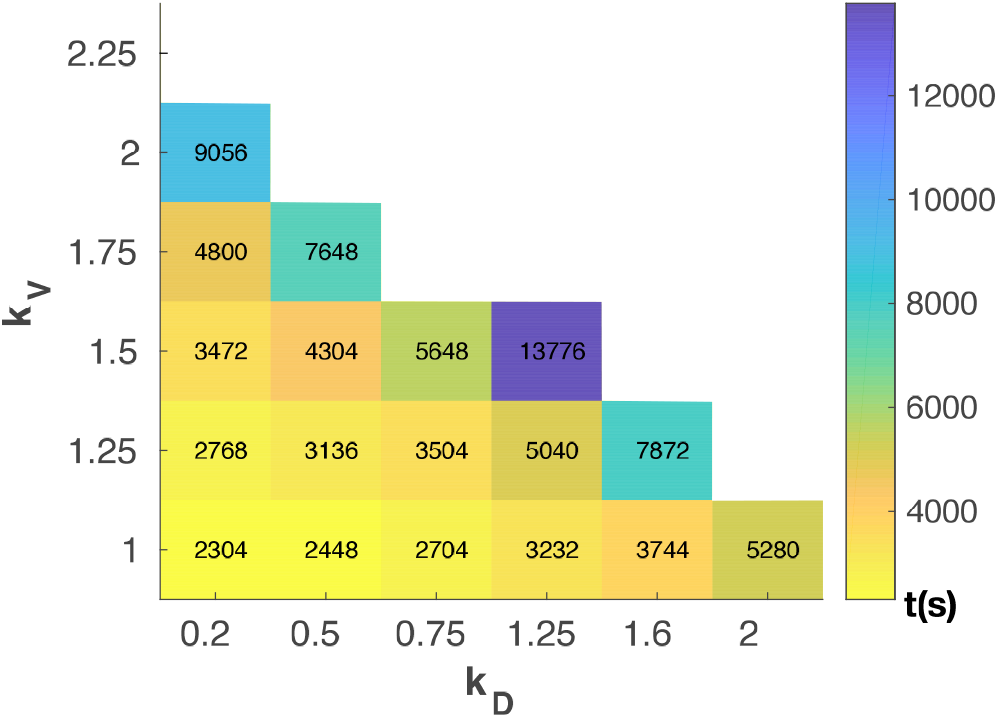
Contour plot of closure time for a range of *k*_*D*_ and *k*_*V*_. The values 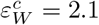 and *δ* = 1.0 have been used. White areas indicate simulations where wound did not close.

**Fig 12.**
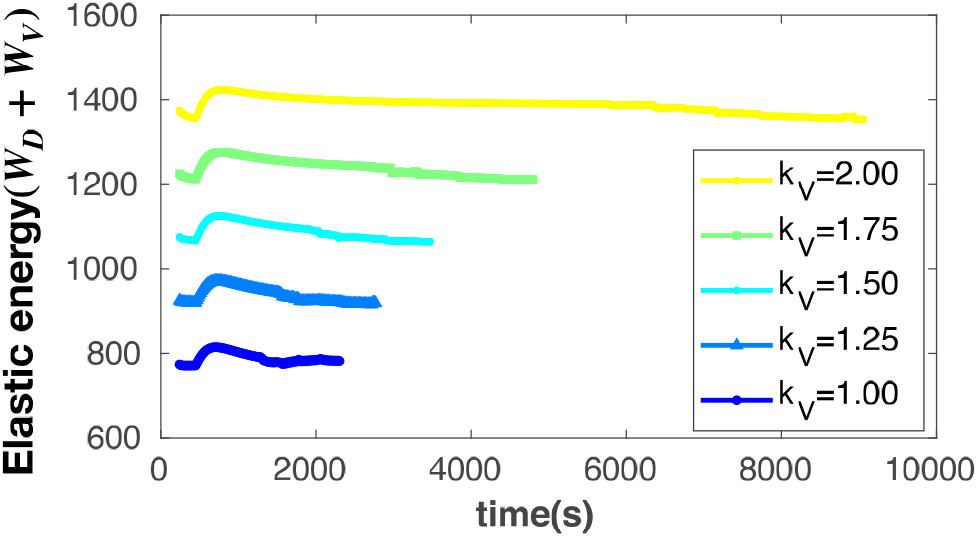
Elastic energy in time for *k*_*D*_ = 0.2 and a range of *k*_*V*_. The values 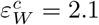 and *δ* = 1.0 have been used.

We also inspect the number of intercalations during the closure process, *N*_*remodel*_. Figure 13 shows the total number in the (*k*_*D*_, *k*_*V*_) and 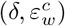 diagrams. We note that remodelling events may be occurring in the bulk tissue rather than specifically at the wound edge, and consequently may not necessarily prevent the formation of rosette-like wounds. For large values of *δ*, most of the intercalations take place at the later stages, while for lower values, they start in earlier stages a proceed at a more constant rate. It is also worth noting that by increasing *δ* at higher values of 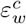, the closure is nevertheless obtained in absence of intercalations. In both cases though, *N*_*remodel*_ has larger variations at lower values of *k*_*V*_ or *k*_*D*_, where larger values of *N*_*remodel*_ are observed.

**Fig 13.**
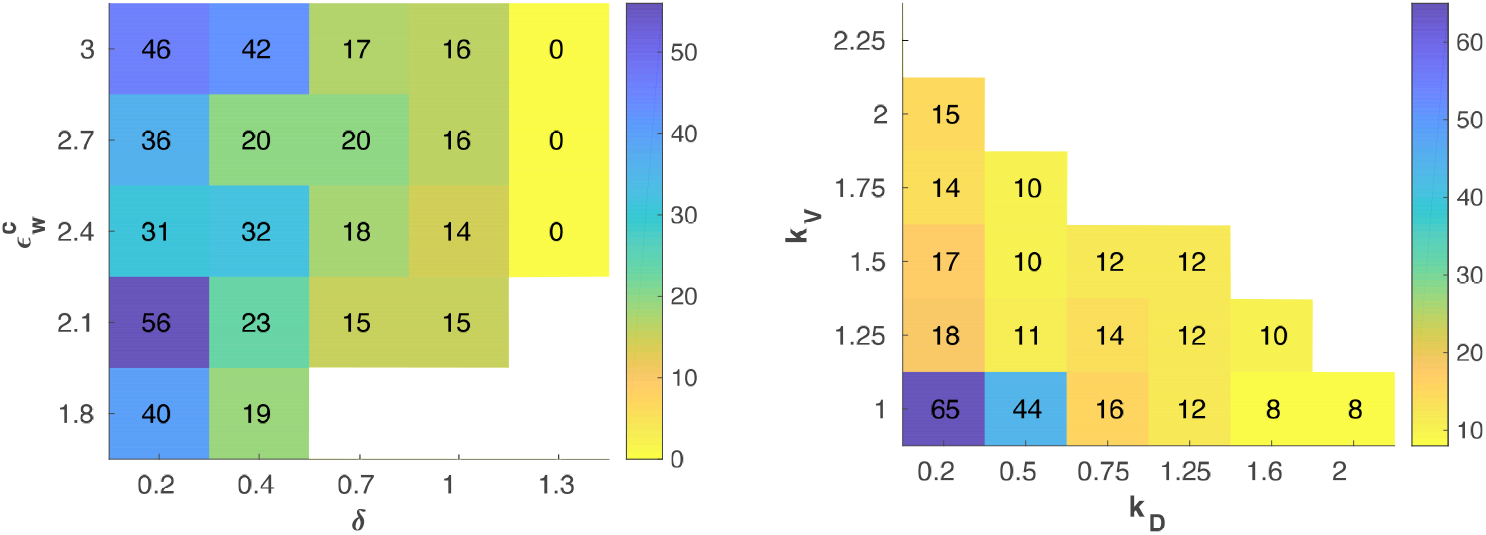
Contour plot of number of remodellings *N*_*remodel*_ in the 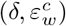 and (*k*_*D*_, *k*_*V*_) planes.

Similar observations are made when varying the contractilities (*ε*_*D*_, *ε*_*V*_) instead of the stiffnesses, as shown in Figure 14. The closure time increases when *ε*_*D*_ or *ε*_*V*_ increases, but vertex contractilities have a more pronounced effect than cell centre ones.

**Fig 14.**
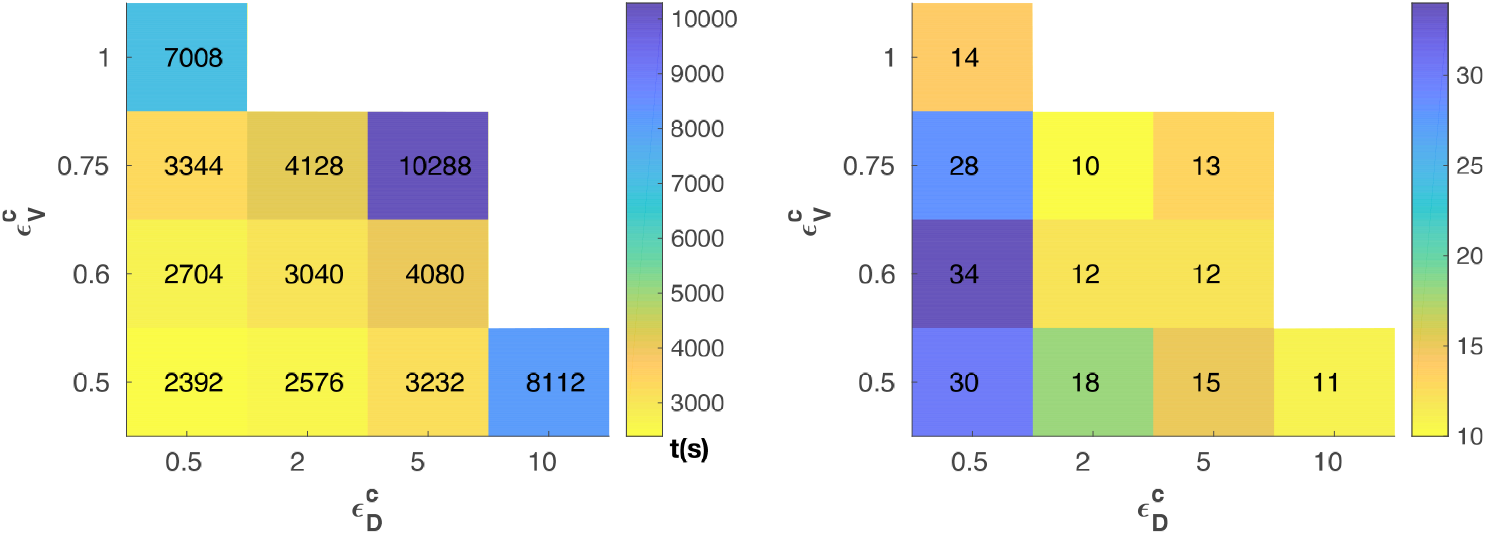
Contour plot of closure time and number of remodelling *N*_*remodel*_ in the 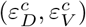 plane. The values *k*_*D*_ = 0.2, *k*_*V*_ = 2.0, 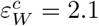 and *δ* = 1.0 have been used. White areas indicate simulations where wound did not close.

The number of remodellings is increased at low values of *ε*_*D*_ and *ε*_*V*_, but do not show in general very large differences at higher values of *ε*_*D*_ and *ε*_*V*_.

The dependence of the closure process on the intercalation rate and tissue fluidity has been recently revealed [4] by varying forces at the cell boundaries. We show here that this dependence also holds true for non-junctional cortically generated forces, with similar but minor effects. Importantly, it has been shown that pulling forces triggered by medial Myosin II flow cannot be discounted [30] in morphogenesis. Our results suggest that such non-junctional cortically generated forces may similarly have an impact on epithelial wound closure.

## Conclusions

We have presented a vertex computational model that is able to distinguish viscoelastic properties at cell junctions and cytoplasmatic regions. The model parameters have been fitted from experimental wound recoil and closure measurements.

After inspection of the energy evolution as a function of stiffnesses in the vertex and nodal network, (*k*_*D*_, *k*_*V*_), wound contractility 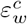, and cell intercalation restriction *δ*, we have identified different mechanisms that can increase closure rate. From our results, an increase in intercalation rate or a reduction in vertex stiffness is more effective than reducing cytoplasmic stiffness or increasing purse cable contractility. The latter induces an important increase in the tissue stored energy confirming the actomyosin purse string is actually dispensable for closure, as also suggested in dorsal closure of *Drosophila* embryo [31].

Our analyses show that the stored energy after closure is slightly higher than just after wounding. This energy gap is higher for increasing actomyosin cable contractiltities, but in all cases lower than the energy of the tissue before wounding. It is yet experimentally unclear though whether the tissue recovers its initial stress and energetic state. Further experiments with rewounding should allow us to quantify whether reepithelialisation rates are altered in recently healed tissues.

Our results confirm that increasing cellular rearrangements can ease wound closure [4]. We further studied the prominent role of junctions instead of other cell mechanical contributions from medial myosin or microtubules. Despite the clear benefit with respect to closure time of increasing intercalation rate and tissue fluidity, it is unlikely that tissues directly aim at reducing the duration of healing. However, compared to other mechanisms such as higher purse string contractility, the higher energy barrier of the latter may prevent the choice of larger values of 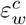 in the material parameter space. Furthermore, fast closure rates have a clear advantage from an evolutionary perspective, which may potentially lead to select the observed mechanisms such as increasing intercalations or lowering junctional stiffness..

### S1 Construction of boundary vertices

According to Figure 15, let us denote by 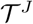 a generic external triangle of the network of cell centres, with external element 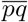 connecting nodes ***x*^*p*^** and ***x*^*q*^** and a corresponding internal vertex ***y*^*J*^**. A primary external vertex 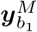 is defined by reflection of ***y*^*J*^** across the line segment 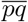, and a similar one is placed outside triangle 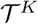. A secondary set of vertices 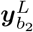 is defined on the arc, 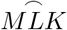, between 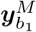 and 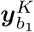. Arc 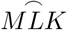 belongs to the circle passing by ***y*^*J*^**, 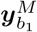 and 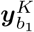. Curve 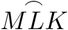 is fragmented equally by the external vertices 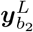. The number of vertices on 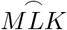 is defined by parameter *α* which is proportional to the length of 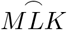.

**Fig 15.**
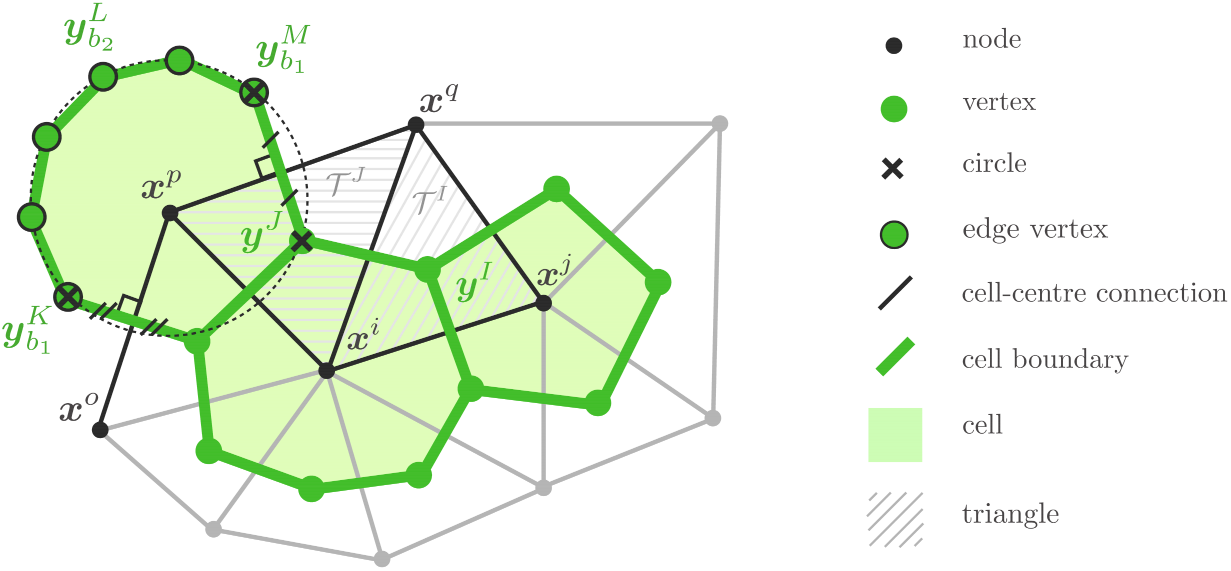
Discretisation of tissue by triangulation of cell centres (***x*^*i*^**), tessellation of the triangulation to obtain cells boundaries (***y*^*I*^**), and definition of cells boundary at the edge of the tissue 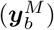.

**Fig 16.**
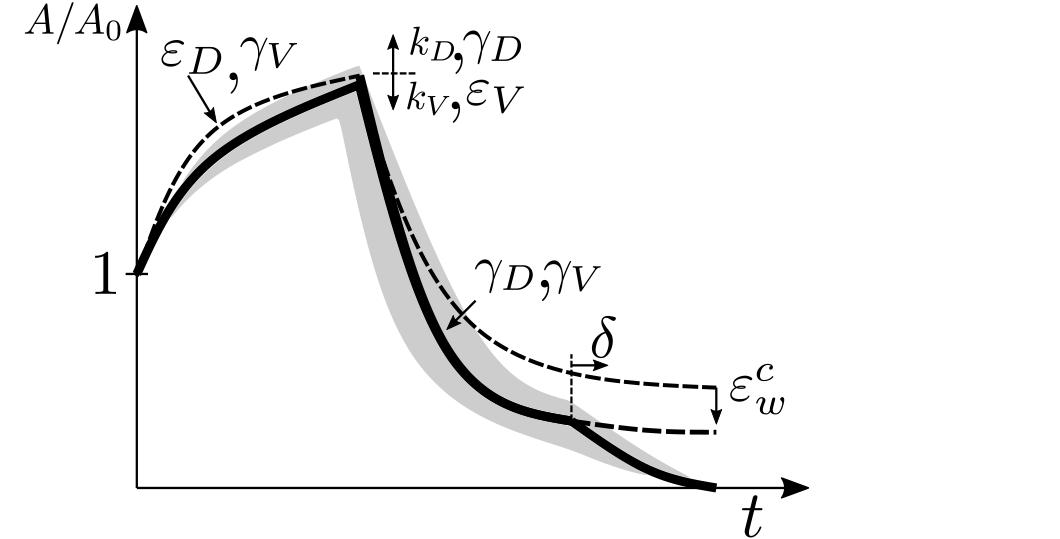
Variation of each background material parameter versus experimental results during the wound recoil. Arrows indicate direction of variation of the curves for increasing values of the parameters.

The radius of the arc, defining the initial position of the vertices is defined from the initial area of the cell. The actual position of the vertices is found through mechanical balance, as described in section Mechanical Equilibrium.

### S2 Fitting of Model Parameters

In this section, we briefly explain the fitting of the material parameters in order to best mimic the wound area evolution during the course of healing in experimental tissues.

Before proceeding to fit material parameters in order to simulate experimental results, we studied the role of each parameter separately.

Wound healing in the *Drosophila* wing disc has been reported to consist of two phases. The first phase corresponds to a nearly immediate fast recoil of the wound edge, while the second phase consists of the gradual shrinkage of the wound area ending in the full closure of the wound. These two phases can be identified in Figure 5.

Since we use the active model to simulate the rheology in both the nodal and vertex networks, representing mechanics of cell bulk and boundary matter respectively, we define the parameters stiffness *k*, remodelling *γ* and contractility *ε*^*c*^ using subscripts *D* and *V* for either of these networks, respectively. Figure 5 shows how variation of each single material parameter can affect the wound recoil during the recoil phase of the wound area evolution.

By obtaining the set of background material parameters which makes the area evolution mimic the experimental results during the first phase, we are able to evaluate the effect of varying wound edge contractility factor 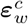 and cell motility *δ* during the second phase of the wound area evolution.

Note that the intercalation *δ* is not considered during the first phase, since due to the short time scale of the area evolution in this phase, no cell intercalation is observed. 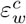 was neither considered during the first phase, since the actomyosin concentration responsible for contractility gain at the wound edge is assumed to occur instantly with a delay respecting the wounding instant. 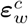 is denoted in units of vertex background contractility 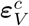. Note also that *k*_*D*_ and *k*_*V*_ have opposite effects on the peak of the area before closure starts.

## Acknowledgments

ARF, PM, JJM have been financially supported by the Spanish Ministry of Science, Innovation and Universities (MICINN) with grant DPI2016-74929-R and by the local government *Generalitat de Catalunya* with grant 2017 SGR 1278. RJT was funded by a Medical Research Council Skills Development Fellowship (MR/N014529/1). YM is funded by a MRC fellowship MR/L009056/1, a UCL Excellence Fellowship, a NSFC International Young Scientist fellowship 31650110472, a Lister Institute Research Prize Fellowship, and EMBO Young Investigator Programme. This work was also supported by MRC funding to the MRC LMCB University Unit at UCL, award code MCU12266B.

